# SFMetaDB: A Comprehensive Annotation of Mouse RNA Splicing Factor RNA-Seq Datasets

**DOI:** 10.1101/177931

**Authors:** Jin Li, Ching-San Tseng, Antonio Federico, Franjo Ivankovic, Yi-Shuian Huang, Alfredo Ciccodicola, Maurice S. Swanson, Peng Yu

## Abstract

Although the number of RNA-Seq datasets deposited publicly has increased over the past few years, incomplete annotation of the associated metadata limits their potential use. Because of the importance of RNA splicing in diseases and biological processes, we constructed a database called SFMetaDB by curating datasets related with RNA splicing factors. Our effort focused on the RNA-Seq datasets in which splicing factors were knocked-down, knocked-out or over-expressed, leading to 75 datasets corresponding to 56 splicing factors. These datasets can be used in differential alternative splicing analysis for the identification of the potential targets of these splicing factors and other functional studies. Surprisingly, only ∼15% of all the splicing factors have been studied by loss- or gain-of-function experiments using RNA-Seq. In particular, splicing factors with domains from a few dominant Pfam domain families have not been studied. This suggests a significant gap that needs to be addressed to fully elucidate the splicing regulatory landscape. Indeed, there are already mouse models available for ∼20 of the unstudied splicing factors, and it can be a fruitful research direction to study these splicing factors in vitro and in vivo using RNA-Seq.

**Database URL:** http://sfmetadb.ece.tamu.edu/

## Introduction

Due to the lack of fully structured metadata, the wide use of the valuable RNA-Seq datasets in public repositories such as ArrayExpress (1) and Gene Expression Omnibus (GEO) (2) may be restricted, despite structured metadata having been used elsewhere for raw data usability (3). For example, ArrayExpress is only a repository of datasets, and the completeness of metadata information relies on dataset submitters. Although submission facilities have been improving, metadata information of many datasets in ArrayExpress is still not well structured (1). To fill this gap, manual curation has been devoted to developing and maintaining metadata databases (4). For example, microarray and RNA-Seq datasets have been curated for the downstream analyses in Expression Atlas (5). We previously launched the RNASeqMetaDB (6) database to facilitate the access of the metadata of public available mouse RNA-Seq datasets. Here, we present a new database, SFMetaDB, as an update with metadata of RNA-Seq datasets related with splicing factors with either loss- or gain-of-function experiments.

RNA splicing is a fundamental biological process in eukaryotes that substantially contributes to the overall protein diversity in a cell. According to GENCODE (Release 25) basic transcript annotation, 19903 human protein-coding genes encode 54896 isoforms by alternative splicing. The importance of alternative splicing is underscored by the distinct biological functions played by splicing isoforms. Recently, the splicing isoform function of a number of genes has been tested experimentally in a variety of biological contexts, including cancer. For example, two isoforms of *CD44*, a widely expressed cell surface marker, have recently been shown to be important in cancer development. The first isoform CD44V6 is required for the migration and generation of metastatic tumors in colorectal cancer stem cells and can initiate the metastatic process (7). The second isoform of *CD44*, CD44V8-10, is an important marker for human gastric cancer and increases tumor initiation in gastric cancer cells (8). Another example is *NUMB*, a gene that is critical for cell fate determination. Two splicing isoforms varying in the length of proline-rich region (PRR), PRR^L^ and PRR^S^, were recently found to have opposite roles in hepatocellular carcinoma (HCC), suggesting that the alternative splicing of *NUMB* can serve as an important biomarker for HCC (9). In particular, PRR^L^ promotes proliferation, migration, invasion and colony formation while PRR^S^ generally works in the opposite way.

Splicing isoforms may also play some critical roles in biological processes other than cancer. For example, *MICU1* is a gene encoding an essential regulator of mitochondrial Ca^2+^ uptake, a process that is critical for energy production in skeletal muscle. Through the inclusion of a micro-exon (<15 bp) of this gene, an alternative splice isoform named MICU1.1 can be generated. It was found that the exclusion of this microexon causes a ∼10x decrease of the Ca^2+^ binding affinity of MICU1 proteins. Therefore, alternative splicing is essential for the sustainability of Ca^2+^ uptake and ATP production of mitochondria, the energy source of skeletal muscle (10). For another example, FANCE is a part of the Fanconi anemia (FA) complex, which functions in DNA interstrand crosslink repair. FANCE plays a critical role to regulate FANCD2, which is required in FANC-BRCA functions. Overexpression of an alternative splicing isoform FANCEΔ4 promotes degradation of FANCD2 and causes dysfunction of DNA repair (11). Furthermore, *VEGF-A* is a gene that functions in angiogenesis, vasculogenesis, and endothelial cell growth. Two alternative splicing isoforms, VEGF-A_xxx_a and VEGF-A_xxx_b, are critical in nociception (12). VEGF-A_xxx_a is increased with nerve injury and promotes nociceptive function. On the contrary, the overexpression of VEGF-A_xxx_b reduces neuropathic pain. In addition, the *Fas*/*CD95* gene is critical in the physiological regulation of programmed cell death. *Fas*/*CD95* has two splicing isoforms with inclusion or exclusion of exon 6, a membrane-bound receptor or a soluble isoform (13). The membrane-bound receptor isoform promotes apoptosis while the soluble isoform inhibits apoptosis.

Alternative splicing is commonly mediated by RNA splicing factors (14). For example, the splicing factor NOVA1 regulates the alternative splicing of a series of genes in pancreatic beta cells, and knockdown of *Nova1* suppresses insulin secretion and promotes apoptosis (15). Moreover, the splicing factor NOVA2 uniquely mediates the alternative splicing of many axon guidance related genes during cortical development (16). As another example, the splicing factor PTBP1 suppresses *Pbx1* exon 7 and the neuronal PBX1A isoform in embryonic stem cells (ESCs) during neuronal development (17).

In this paper, we describe our recent effort in curating the metadata of RNA-Seq datasets from ArrayExpress and GEO, which were derived from studies using cell or animal models with a specific splicing factor being knocked-out, knocked-down, or overexpressed. We further launched SFMetaDB to facilitate access to the metadata of these datasets and share them with the biomedical community.

## Results and Discussion

The launch of SFMetaDB focuses on RNA-Seq datasets with perturbed splicing factors. Users can query a given splicing factor to identify the relevant datasets. A use case for MBNL splicing factors is shown as follows. MBNL1 is an important RNA splicing factor (18), thus we use MBNL1 to demonstrate the usage of SFMetaDB, which confirms the advantage of SFMetaDB over ArrayExpress. As shown in Figure 1a, a query of MBNL1 on SFMetaDB returns the accurate datasets related with *Mbnl1* loss- or gain-of-function experiments. Figure 1a shows that five datasets could be used for the alternative splicing analysis for MBNL1, and the potential targets of MBNL1 can be concluded from the datasets. For example, the dataset GSE39911 (i.e. E-GEOD-39911) includes biological replicates of various tissues, such as brain, heart and muscle, from *Mbnl1-*knockout mice and *Mbnl1-*knockdown C2C12 mouse myoblasts (Figure 1b).

**Figure 1.**
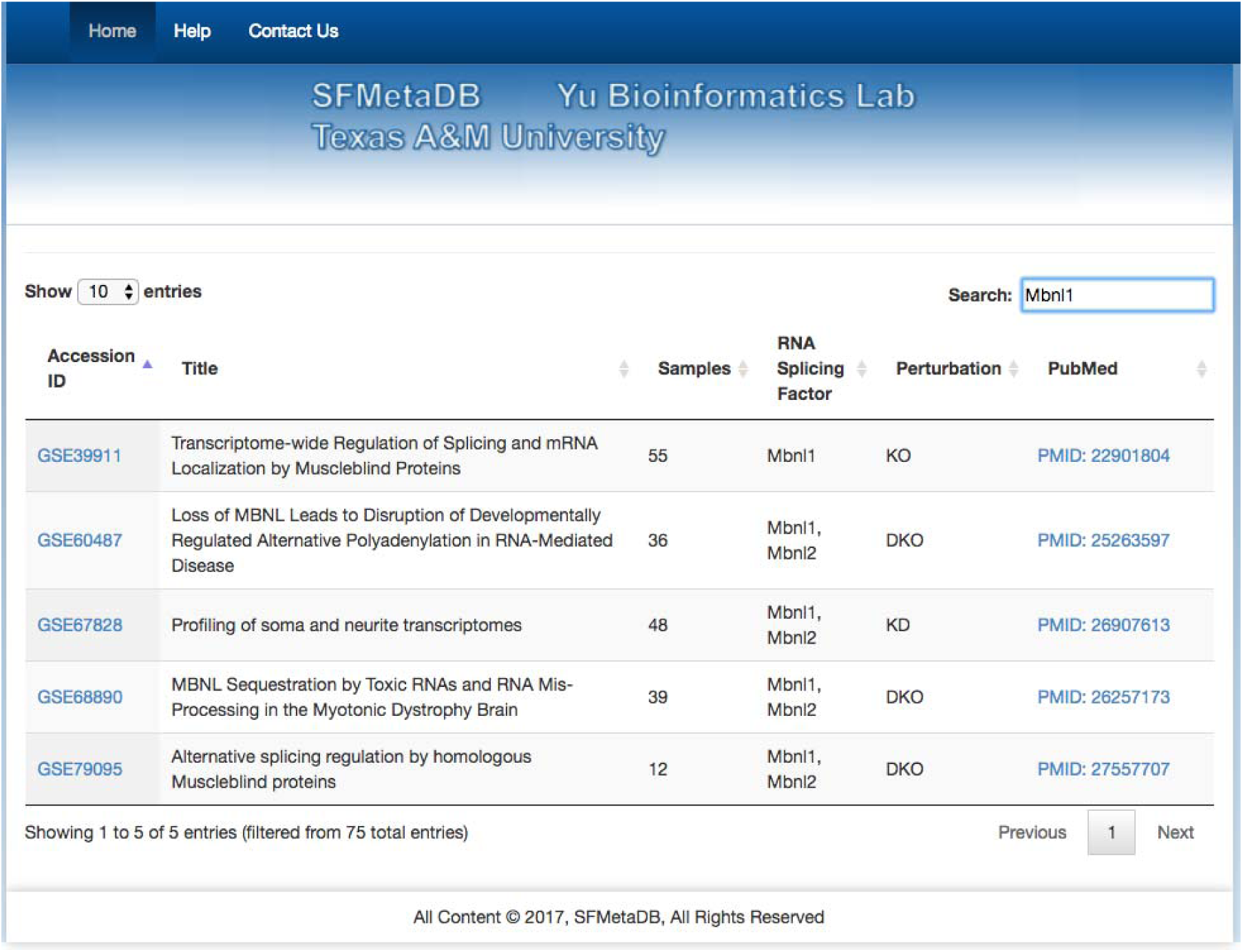
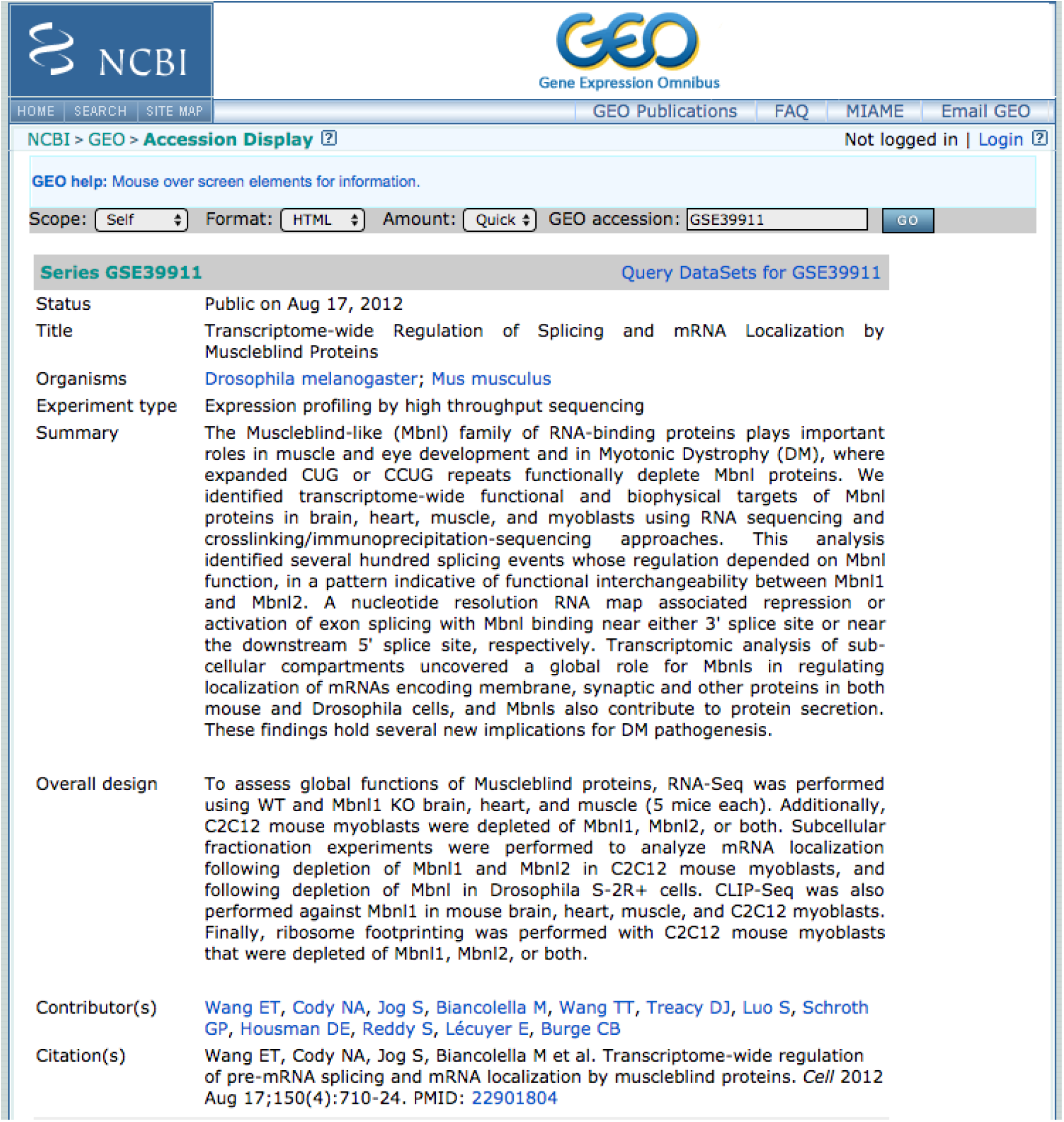
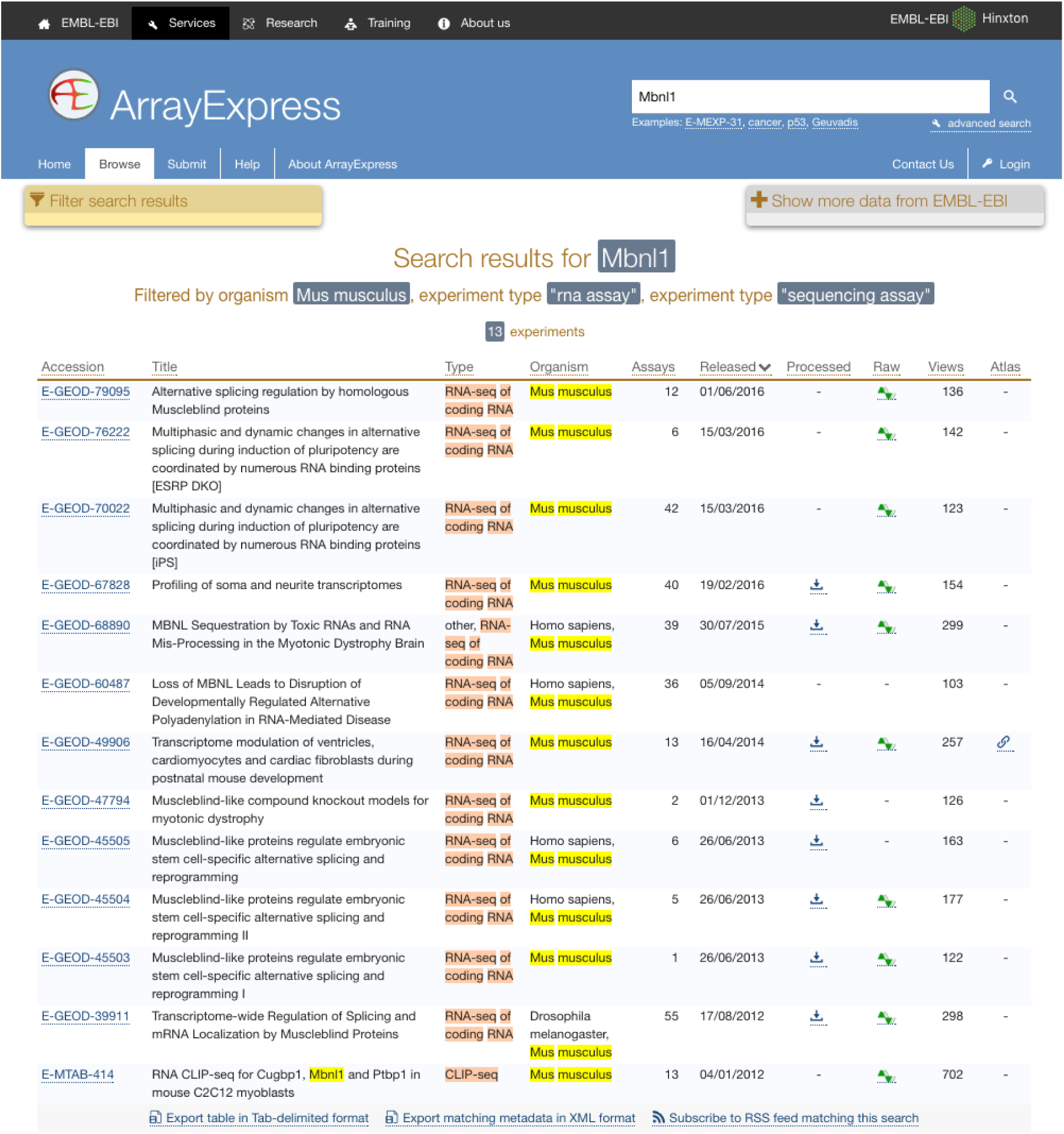
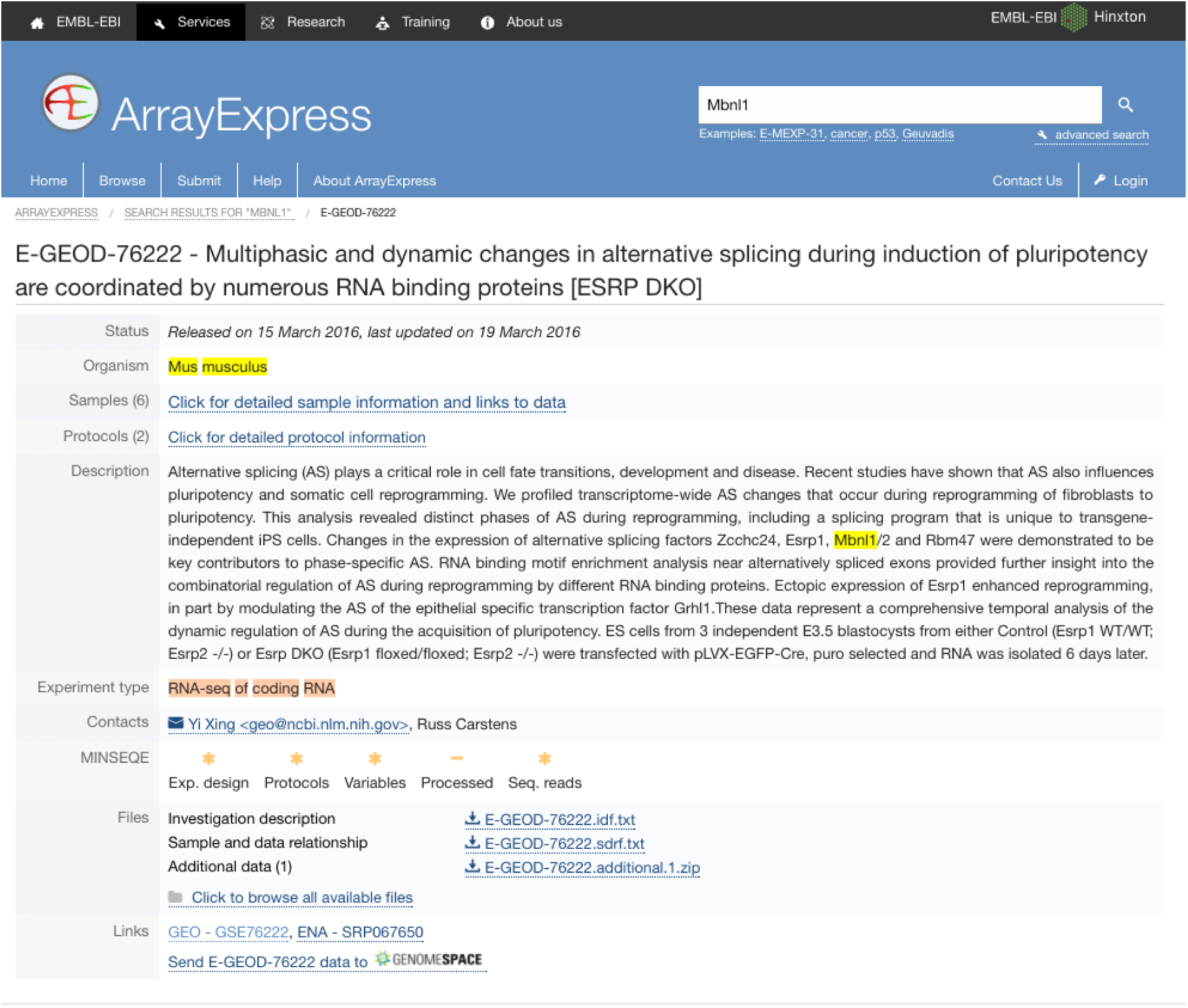
A use case of SFMetaDB for the splicing factor Mbnl1. We showed a use case of the splicing factor Mbnl1 to demonstrate the advantage of SFMetaDB over ArrayExpress. By using the same keyword, Mbnl1, SFMetaDB returned five accurate datasets that can be used for the downstream alternative splicing analyses. On the contrary, ArrayExpress returned thirteen datasets with eight that could not be used for the downstream alternative splicing analyses for Mbnl1. (a) The result page in SFMetaDB of the query Mbnl1. (b) The description page of the dataset GSE39911 in GEO. (c) The result page in ArrayExpress of the query Mbnl1. (d) The description page of the dataset E-GEOD-76222 in ArrayExpress.

However, as shown in Figure 1c, ArrayExpress returned a total of 13 mouse RNA-Seq datasets with the query Mbnl1, and eight of them were not from *Mbnl1* gain- or loss- of function experiments. Therefore, these datasets were eliminated in SFMetaDB. For example, the dataset E-GEOD-76222 is retrieved by ArrayExpress because of the appearance of Mbnl1 in its description, “Changes in the expression of alternative splicing factors Zcchc24, Esrp1, Mbnl1/2 and Rbm47 were demonstrated to be key contributors to phase-specific AS.” However, this dataset is about an *ESRP* knock-out, thus it is not suitable for MBNL1 related alternative splicing analysis (Figure 1d). The rest of eight retrieved datasets were considered not appropriate for RNA splicing analysis of MBNL1 by our manual curation of metadata information. In summary, no irrelevant datasets of a given splicing factor are shown in SFMetaDB, and SFMetaDB returned more specific results than ArrayExpress.

Guided by SFMetaDB, users can perform potential target identification for a specific splicing factor. In addition, by integrating multiple datasets curated on SFMetaDB, users can form a more comprehensive view on how a splicing event is regulated across different biological contexts. As another use case, we show below a Pfam domain analysis among splicing factors (See Materials and methods).

Only ∼15% of known splicing factors have been studied with loss- or gain-of-function RNA-Seq experiments. Because splicing factors sharing similar domains tend to regulate common splicing targets, we determined what additional splicing factors may be prioritized for study by investigating the domain structures of the splicing factors using UniProt (19). Among the 353 splicing factors, 299 of them contained one or multiple conservative domains. Of these 299 splicing factors, 190 have a single domain that belongs to a Pfam domain family, and the rest have domains that belong to more than one Pfam domain family.

RNA splicing factors have highly conserved functional domains, and some domains are dominant among all the splicing factors. In Figure 2, the domain families are ranked by their number of occurrences in all the splicing factors. Pfam family PF00076 (RNA recognition motif) is the most dominant, and the splicing factors with domains from this family are relatively well-studied (25 over the total 87). Splicing factors from five additional Pfam families are fairly well-studied (≥ 3 splicing factors annotated), consisting of PF00271(Helicase conserved C-terminal domain), PF00270(DEAD/DEAH box helicase), PF00013(KH domain), PF00642 (Zinc finger C-x8-C-x5-C-x3-H type) and PF12414 (Calcitonin gene-related peptide regulator C terminal). However, three highly dominant families are not. Specifically, none of the 17 splicing factors with the Pfam family PF01423 (LSM domain) (Figure 2) have been studied yet (20), and these splicing factors provide feasible candidates for future studies. For example, the splicing factor SNRPN has two mouse models from the International Mouse Strain Resource (IMSR) (21) that can be used for splicing analysis. In fact, twenty-five unstudied splicing factors (**Table S1**) have been identified with more than one mouse model from IMSR. Therefore, splicing factors that are non-homologous with already studied ones constitute promising candidates for comprehensive studies of splicing regulation.

**Figure 2.**
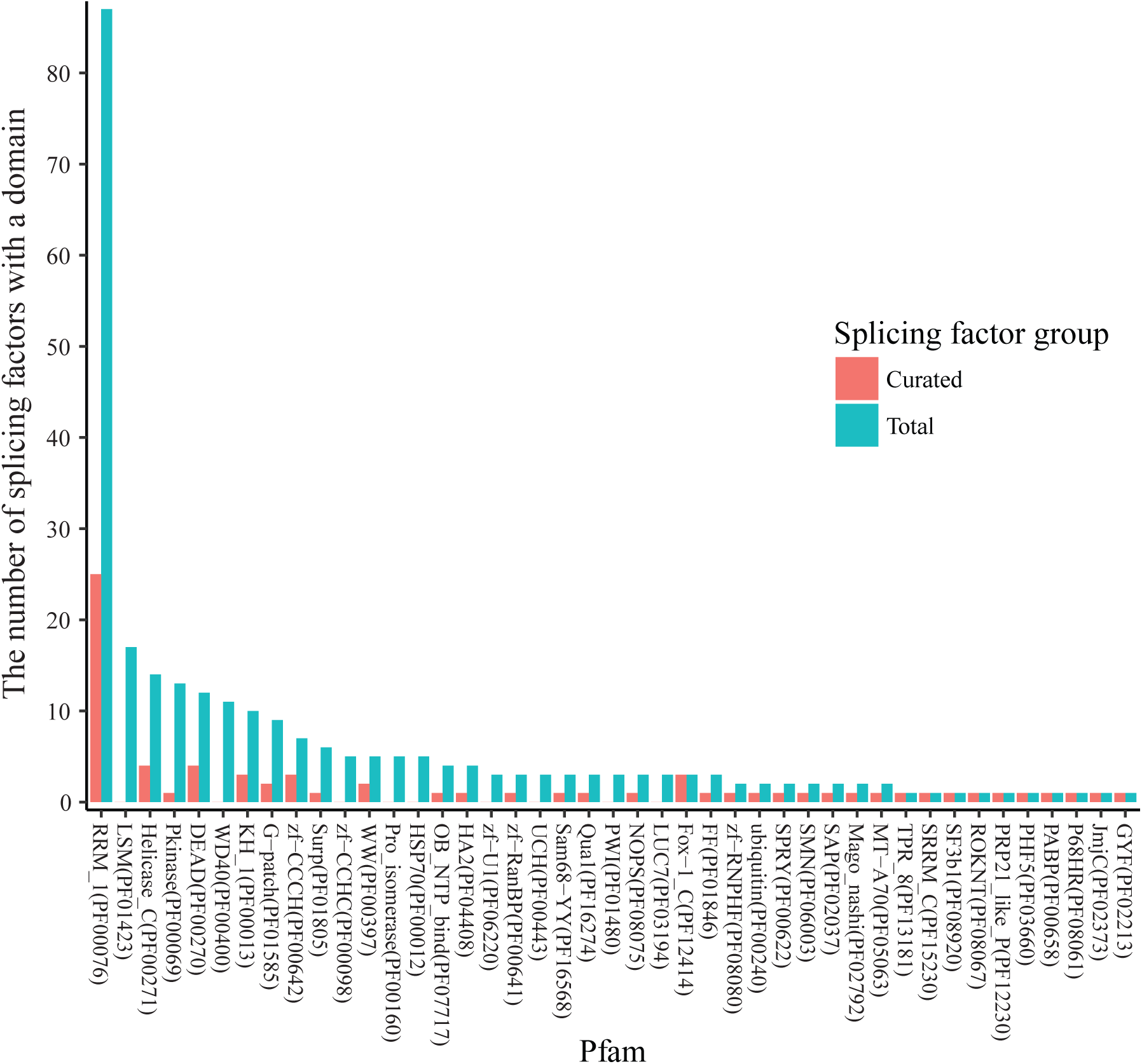
The occurrence of Pfam domain families in splicing factors. The known RNA splicing factors are annotated in UniProt according to the Pfam domain families of the protein domains found in these factors. A splicing factor may have multiple domains that belong to multiple Pfam families, and a Pfam domain family may contain domains in multiple splicing factors. The Pfam annotations were retrieved for each of 353 splicing factors, and the number of splicing factors was calculated for each of the Pfam families. For the 56 splicing factors that have curated datasets in SFMetaDB, the number of splicing factors was also calculated for the associated Pfam families. In the dodged barplots, the Pfam domain families are ranked by the number of the splicing factors which contain domains in the given families. Of the total 217 Pfam domain families annotated in UniProt, 26 Pfam domain families have ≥ 3 splicing factors annotated. The Pfam domain family with the most number of splicing factors is Pfam RRM_1 (PF00076). It contains 87 splicing factors, and 25 of these splicing factors have been studied according to our curation results. However, the splicing factors in the rest of the Pfam domain families have brought relatively less attention in RNA-Seq analysis, and they may be promising candidates for future studies.

## Materials and methods

### RNA-Seq dataset curation and SFMetaDB web server deployment

We extracted 353 RNA splicing factors annotated in Gene Ontology (GO) (accession GO:0008380) (22) and Kyoto Encyclopedia of Genes and Genomes (KEGG) (entry mmu03040) (23) for mice. Then, we queried ArrayExpress (1) and GEO (2) using the official symbol of each splicing factor to search for related mouse RNA-Seq datasets and obtained a total of 214 datasets. Note that due to the limitation of the search function in ArrayExpress and GEO, many of these datasets were not directly relevant to the manipulation of these splicing factors despite that the symbols were mentioned in the metadata of these datasets. We chose to manually curate each dataset, providing a total of 75 datasets that have biological replications in which at least one splicing factor was knocked-out, knocked-down or overexpressed (along with the corresponding wild types/controls) (**Table S1**). Because some splicing factors were studied in more than one dataset, a total of 56 splicing factors were found (**Table S1**).

To facilitate the access to these datasets, we launched the database SFMetaDB (http://sfmetadb.ece.tamu.edu/). When datasets were deposited in GEO, ArrayExpress imported the most metadata information from GEO, and the ArrayExpress description contained the link to the GEO webpage. Therefore, SFMetaDB used GEO accession ID if possible. The web server of SFMetaDB is freely available, and it presents the Accession ID, description, the number of samples, associated curated splicing factors, perturbation and PubMed references of each RNA-Seq dataset.

### Domain structures analysis in RNA splicing factors

The domain structures of the RNA splicing factors may guide us to identify the candidate splicing factors for future studies. Known RNA splicing factors are retrieved from GO term (GO:0008380) using R package GO.db (22) and KEGG pathway (entry mmu03040). UniProt annotates the conservative Pfam domain families for the canonical sequences of the splicing factors (19). From these domain annotations, we calculate the numbers of the splicing factors in Pfam domain families. Figure 2 plots the dodged barplots of the number of splicing factors in Pfam domain families using curated splicing factors and the total splicing factors. By comparing the domain families of the splicing factors with RNA-Seq datasets to the families of all the splicing factors, the splicing factors in not well-studied domain families can be the promising candidates for future RNA-Seq studies.

## Acknowledgements

The authors thank Zhengyu Guo for his contribution to SFMetaDB.

## Supplementary material

**Table S1 Pfam domain families and IMSR mouse models of splicing factors** Splicing factors were annotated with Pfam domain families which contained protein domains from these splicing factors. For each Pfam domain family, the Pfam Accession, the number of splicing factors annotated, the symbols of the splicing factors annotated, Pfam family ID, and Pfam family description are listed. The available mouse models for the splicing factors were searched on the International Mouse Strain Resource (IMSR). For each splicing factor, the Strain/Stock of the mouse models, the associated allele ID, allele symbol, and MGI gene ID are listed. Splicing factors that are not currently well-studied but have mouse models may be ideal candidates for future studies.

